# High-Precision Lighting for Plants: Laser Diodes Outperform LEDs in Photosynthesis and Plant Growth

**DOI:** 10.1101/2025.02.24.640008

**Authors:** Lie Li, Ryusei Sugita, Kampei Yamaguchi, Hiroyuki Togawa, Ichiro Terashima, Wataru Yamori

## Abstract

The optimization of plant productivity in indoor horticulture relies heavily on artificial light systems, which serve as the primary light source for plant growth. Although light-emitting diodes (LEDs) have been extensively studied in recent decades, there is limited research on laser diodes (LDs). LDs offer several advantages such as adjustable light wavebands, remote illumination that minimizes heat accumulation near plants, and enhanced energy efficiency. This study investigated the impact of red LD light on plant photosynthesis and growth, exploring its potential applications in indoor horticulture. The research examined the gas exchange of tobacco plants (*Nicotiana tabacum* L. cv. Wisconsin-38) under six red LED and LD light sources with varying spectral characteristics. Two specific light sources were selected for further study: LED 664 (emission peak at 664 nm, waveband of 625~678 nm) and LD 660 (emission peak at 660 nm, waveband of 657~664 nm) as they demonstrated the greatest gas exchange efficiency among the tested LED and LD light sources. These two light sources were then evaluated for their effects on photochemical efficiency, carbohydrate accumulation and plant growth. The present study showed that compared with LED 664, LD 660 significantly increased Y(II), qL, and starch accumulation in tobacco leaves. Additionally, after 12 d of continuous irradiation with LD 660, both tobacco and Arabidopsis plants exhibited increased photosynthetic capacity, and all three plant species showed increased shoot dry weights and leaf areas compared with those under LED 664. These findings suggest that LDs present significant advantages over LEDs for indoor plant production.

## 1. Introduction

The global population is projected to reach 9.7 billion by 2050, with 70% living in urban areas, which increases food demand while decreasing the agricultural workforce (UN, 2018; UN, 2022). Simultaneously, extreme climate events and geopolitical conflicts are exacerbating the global food crisis (Binns et al., 2021; Vogel et al., 2019; Behnassi et al., 2022; Qu et al., 2023). Owing to its advantages over traditional farming methods, indoor horticulture can play a pivotal role in sustainable food production if key challenges such as energy, labor, and production unit economics are solved. If these challenges can be overcome, indoor horticulture offers benefits such as year-round plant production, reduced labor requirements, no geographical or natural restrictions, high land use efficiency, and precise environmental control (Kozai et al., 2013; Bantis et al., 2018). Identifying an optimal artificial light source is essential, as artificial lighting systems often represent a major capital expenditure due to their high energy consumption (Levine et al., 2024).

Conventional artificial light sources, such as high-pressure sodium (HPS), metal halide (MH), incandescent (INC) lamps, and fluorescent tubes (FTs), have been widely used in greenhouses and plant growth chambers. However, these options are limited as horticultural lighting solutions because their broad emission spectra and low electricity-to-light energy conversion efficiencies limit their effectiveness. Since the 1990s, light-emitting diodes (LEDs) have been increasingly adopted for plant growth, offering advantages such as adjustable spectral composition, lower heat emission, and longer lifespan (Bula et al., 1991; Massa et al., 2008; Bantis et al., 2018; Van Delden et al., 2021).

Compared with traditional light sources and LEDs, laser diodes (LDs) offer distinct advantages such as single-wavelength coherent light, compact size and lightweight design, remote light delivery, less heat generation, low power consumption and high efficiency.

“Laser” is an acronym for “light amplification by stimulated emission of radiation.” In the stimulated emission process, an incoming photon of a specific frequency interacts with an excited atom, electron, or molecule, prompting a transition to a lower energy level. The released energy then transfers to the electromagnetic field, generating a new photon with the identical frequency, polarization, and direction as the incident photon. Consequently, LD light is a single-wavelength, phase-aligned and very intense light (Fain and Milonni, 1987; Klimek-Kopyra et al., 2021). Over recent decades, numerous studies have focused primarily on how light wavelengths affect plant growth and development (Ouzounis et al., 2015), leading to the conclusion that red (600~700 nm) and blue (400~500 nm) light are optimal wavelengths for promoting plant production in indoor horticulture (Van Delden et al., 2021). However, how variation in waveband width impacts photosynthesis and growth remains unclear. While LEDs typically emit light across relatively broad wavebands of approximately 50 nm, LDs emit light within an extremely narrow, adjustable wavelength, typically less than 10 nm. This narrow spectral output allows LDs to closely match the peak light absorption of chlorophyll. Because the absorption efficiency of chlorophyll decreases as the wavelength deviates from its peak, LDs are theoretically more effective at promoting plant growth (Taiz and Zeiger, 2015).

LDs also have the outstanding feature of being compact and lightweight. The laser chip itself typically measures between 100 µm and 3 mm, and the packaging in which the chip is embedded is similarly small, often only a few millimeters in size. LDs can efficiently transmit light over long distances via optical fibers and expand the irradiation range using diffusers, reducing heat generation near plants and conserving space (Hitz et al., 2012; Ooi et al., 2016; Murase, 2015). Furthermore, LDs have the potential to achieve high electricity-to-light conversion efficiency, even at elevated input power densities (Wierer et al., 2013; 2014). Consequently, LDs are anticipated to increase crop yields while promoting environmentally friendly and energy-efficient practices in plant production.

Various plant species have demonstrated increased seed germination rates and enhanced resistance to certain biotic and abiotic stresses following laser seed irradiation (Krawiec et al., 2012; Klimek-Kopyra et al., 2020; Nadimi et al., 2022, Metwally et al., 2014; Gao et al., 2015, 2018; Siyami et al., 2018; Ali et al., 2020). Although these studies provide valuable insights, most have utilized different types of laser lights, such as solid-state lasers, gas lasers and diode lasers, primarily as short-term biostimulants (Klimek-Kopyra et al., 2021). Therefore, further research is essential for the full application of laser light in plant production as a primary artificial light source in indoor horticulture.

The purpose of the present research was to examine both the instantaneous and the acclimation effects of LD light on the photosynthetic performance of three plant species: Arabidopsis, tobacco and lettuce. We utilized six red light sources, four LDs and two LEDs, with different emission peaks and bandwidths.

## 2. Materials and methods

We conducted two experiments using tobacco (*Nicotiana tabacum* L. ‘Wisconsin-38’), *Arabidopsis thaliana* (L.) Heynh ‘Col. 0’, and lettuce (*Lactuca sativa* L. ‘red fire’).

### 2.1 Experiment 1. Effects of different red light spectra on photosynthesis

#### 2.1.1 Plant materials and cultivation conditions

Tobacco seeds were sown in a 1:1 mixture of vermiculite and peat with initial nutrition (Metro-Mix 350J, Hyponex, Japan) in black plastic trays (24.0 cm^3^, 25 x 15 mm, and 45 mm in height). After germination, the seedlings were thinned to one plant per pot. The plants were then grown in a growth chamber (LPH-411SPC, Nippon Medical & Chemical Instruments, Japan) with a 10 h light/14 h dark cycle at an air temperature of 25/22 °C and a relative humidity of 60 ± 5%. Light was provided by white fluorescent tubes at a photosynthetic photon flux density (PPFD) of 120 μmol m^−2^ s^−1^ at the plant level. The CO_2_ concentration was 400 μmol mol^−1^. After 25 d of seedling growth, four fully expanded young leaves from different plants were selected to measure gas exchange, and another set of four different plants was used to measure chlorophyll fluorescence parameters.

#### 2.1.2. Gas exchange measurements

The net photosynthetic rate and stomatal conductance were measured in fully expanded tobacco leaves using a portable gas exchange instrument equipped with a transparent leaf chamber (LI-6400XT and 6400-8, LI-COR Biosciences, Lincoln, NE, USA) (Joshi et al., 2017). These measurements were performed under leaf chamber conditions set at an air temperature of 24 °C, a CO_2_ concentration of 400 μmol mol^−1^, and a leaf-to-air vapor pressure deficit (VPD) of 0.7–1.0 kP. The light sources included an LED 629 (peak wavelength 629 nm, waveband of 597~645 nm) (ISLM-150X150-RB, CCS Inc., Japan), an LED 664 (peak wavelength 664 nm, waveband of 625~678 nm) (3LH-100DPS, Nippon Medical & Chemical Instruments, Japan), an LD 635 (peak wavelength 635 nm, waveband of 631~638 nm) (HL63283HD, Ushio Inc., Japan), an LD 660 (peak wavelength 660 nm, waveband of 657~664 nm) (HL65213HD, Ushio Inc., Japan), an LD 673 (peak wavelength 673 nm, waveband of 669~677 nm) (HL67203HD, Ushio Inc., Japan), and an LD 690 (peak wavelength 690 nm, waveband of 685~693 nm) (HL69203HD, Ushio Inc., Japan), each providing a PPFD of 150 μmol m□² s□¹. Each of these six light sources was applied to each of the four leaves in a random sequence, and the data were recorded once the values stabilized. The spectra of these light sources were determined using a spectroradiometer (LA-105, Nippon Medical & Chemical Instruments Co., Ltd., Japan) and are shown in Figure 1. The intrinsic water use efficiency (WUE_i_, mmol CO_2_ mol^−1^ H_2_O) was calculated by dividing the net photosynthetic rate by the stomatal conductance.

**Figure 1:**
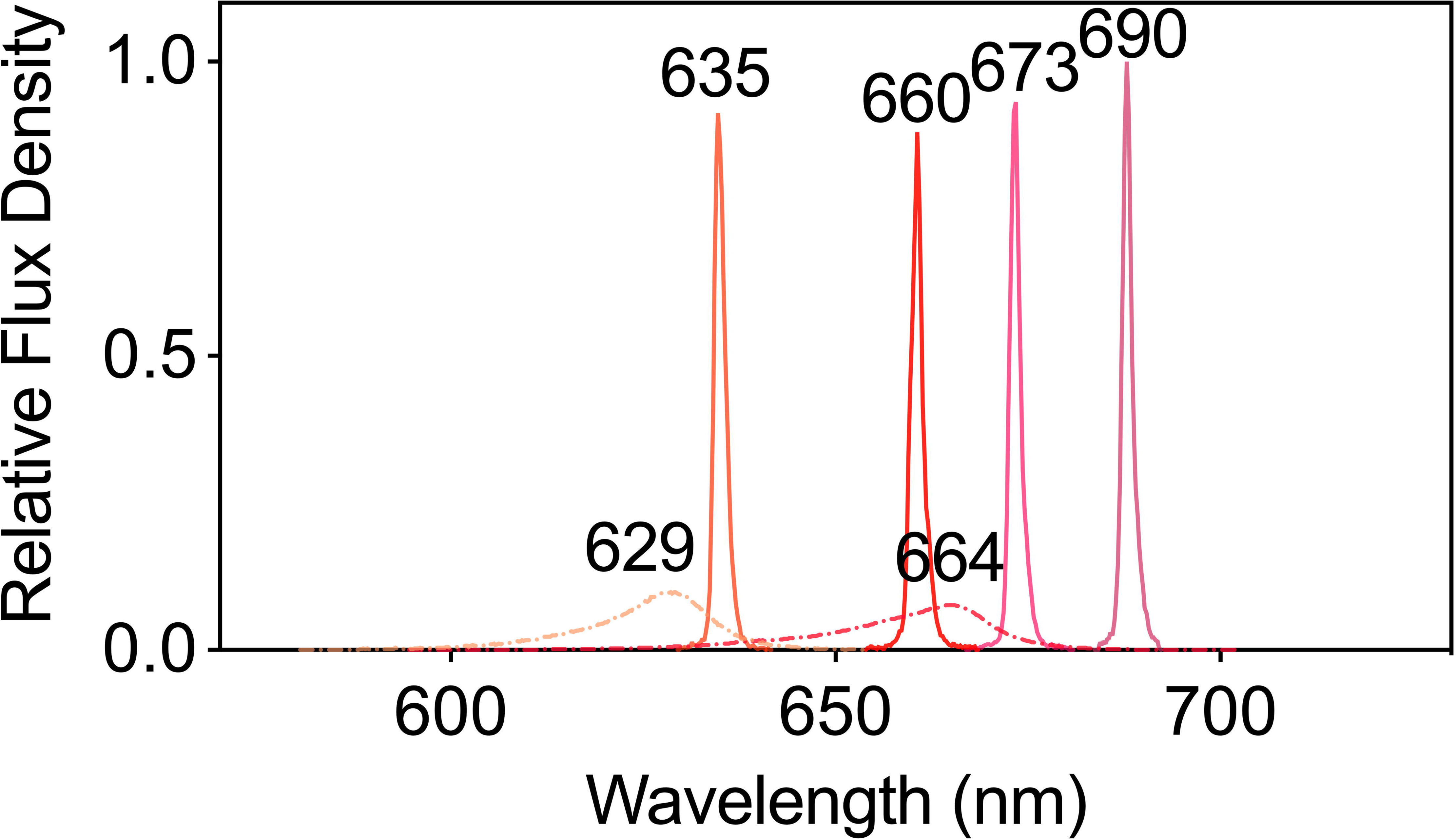
The spectra of plants subjected to six different LED and LD lights were used to test their photosynthetic performance. The dashed lines represent LED lights, whereas the solid lines represents LD lights.

#### 2.1.3. Chlorophyll fluorescence analysis

We used LED 664 and LD 660 light sources to investigate the photochemical efficiency and redox state of PS□, as these light sources supported the highest photosynthetic rates reported in Section 2.1.2. The chlorophyll fluorescence parameters of fully expanded young leaves of tobacco plants were measured using a pulse amplitude modulation fluorometer (Junior-PAM, Heinz Walz GmbH, Germany) combined with fluorescence quenching analysis using saturation pulses. After the plant leaves were dark-adapted for approximately 30 min, the values of minimal fluorescence (F_o_) and maximal fluorescence (F_m_) were measured. Subsequently, either LED 664 or LD 660 at the same PPFD of 150 μmol m□² s□¹ was applied to each of the four leaves in a random sequence. Once a steady-state condition was reached, the steady-state fluorescence (F_s_’) and maximum fluorescence in light (F_m_’) were determined. The minimum fluorescence in light (F_o_’) was calculated as (Oxborugh and Baker, 1997):

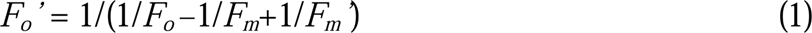

The quantum efficiency of PSII electron transport in light (Y(□)), nonphotochemical quenching (NPQ), and the estimated fraction of open PSII centers (qL) were calculated as follows (Hormann et al., 1994; Murchie and Lawson, 2013):

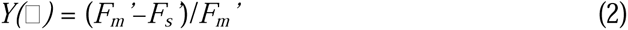

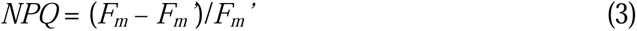

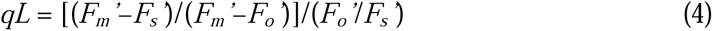

#### 2.1.4. Carbohydrate accumulation

After 25 d of growth, as described in Section 2.1.1, the tobacco plants were illuminated with either of the light sources at a PPFD of 150 μmol m^−2^ s^−1^ for 8 h. The illumination was started at the same time each day. Leaves were detached from each plant and scanned with a scanner (CanoScan LiDE 220, Canon Inc., Japan), and their areas were measured using image analysis software (ImageJ 13.0.6, Wayne Rasband and contributors, National Institutes of Health, USA). The samples were placed into tin foil bags, quick-frozen in liquid nitrogen, and stored at –80 °C until use.

The plant tissues were ground into powder in liquid nitrogen with a mortar and pestle. Each powdered sample was placed in a 2 mL tube. One milliliter of 80% EtOH was added, followed by vigorous vortexing. The samples were then heated to 80 °C for 10 min in a block incubator to extract soluble sugars. The samples were subsequently centrifuged at 13,470 × *g* for 5 min at room temperature using a high-speed microcentrifuge (MX-307, Tomy Digital Biology Co., Ltd., Japan). The supernatant and pellet were separated for the determination of soluble sugars and starch, respectively. Starch degradation by glucoamylase and sucrose cleavage by invertase (β-fructosidase) were conducted according to the methods of Bergmeyer and Bernt (1974) and Schmidt (1961), respectively. The spectroscopic assays at 340 nm were conducted using a microplate reader (Synergy H1, BioTek Instruments, Inc., USA). The contents of soluble sugars and starch were then calculated based on these assays.

### 2.2. Experiment 2. Photosynthesis and plant growth under LED 664 and LD 660

#### 2.2.1. Plant materials, light treatments, and cultivation conditions

We selected LED 664 and LD 660 for further experiments. Tobacco seeds were sown as described in Section 2.1.1. Arabidopsis seeds were sown in the substrate mix in plastic boxes (500 cm^3^, 95 x 80 mm and 65 mm in height), and the seedlings were thinned to three per box. Lettuce seeds were sown in rockwool cubes (40 x 30 mm and 40 mm in height), which were hydrated with distilled water and irrigated weekly with a 1/1000 strength nutrient solution with an NPK ratio of 6:10:5 (Hyponex, Hyponex, Japan). Seedlings of tobacco, Arabidopsis, and lettuce were grown under the conditions described in Section 2.1.1. When the plants developed three fully expanded true leaves (after 25 d for tobacco, 24~28 d for Arabidopsis, and 10 d for lettuce), we began continuous 24-h irradiation with LED 664 or LD 660 at a PPFD of 150 μmol m^−2^ s^−1^ for 12 d. The air temperature, relative humidity and ambient CO_2_ concentration were maintained at 24 ± 1 °C, 60 ± 5%, and 400 μmol mol^−1^, respectively. After 12 d of growth, the photosynthetic characteristics and growth indices were quantified for four plants per species and treatment. Pictures of the top surfaces of the plants were taken, and from these images, one representative picture for each plant species and each treatment was selected and is shown in Figure 6A.

#### 2.2.2 Analysis of yield photon flux density (YPFD) and phytochrome photoequilibria (PPE)

The spectral photon distribution (SPD) of each treatment was determined using a spectroradiometer (LA-105, Nippon Medical & Chemical Instruments Co., Ltd., Japan), and the spectra are shown in Figure 1.

The yield photon flux density (YPFD) was calculated as follows:

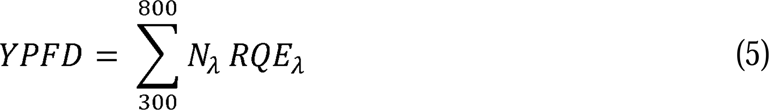

where N_λ_ is the incident flux density of the SPD at wavelength λ, and RQE is the relative quantum efficiency at wavelength λ based on McCree (1972) and Sager et al. (1988).

The phytochrome photoequilibria (PPE) was calculated as follows:

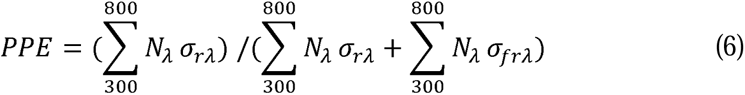

where σ_rλ_ is the red absorbing state of phytochrome photochemical cross-sections at wavelength λ, and σ_frλ_ is the far-red absorbing state.

#### 2.2.3. Chlorophyll fluorescence analysis

The chlorophyll fluorescence parameters and canopy images of tobacco, Arabidopsis, and lettuce were measured using an imaging pulse amplitude modulation fluorometer equipped with a camera (Imaging PAM MAXI version, Heinz Walz GmbH, Germany) (Shimadzu et al., 2019). The actinic light source was blue light with a peak wavelength of 450 nm. After the plants were dark-adapted for approximately 30 min, the values of minimal fluorescence (F_o_) and maximal fluorescence (F_m_) were measured to determine the maximum quantum efficiency of PS □ (F_v_/F_m_), which is equivalent to the Y(□) value at a PPFD of 0. The plants were subsequently illuminated at PPFDs of 37, 98, 189, 311, and 467 μmol m^−2^ s^−1^. After each PPFD was applied for 5 min, F_s_’ and F_m_’ were recorded. Y(□) and NPQ were calculated as described in Section 2.1.3. The F_v_/F_m_ was calculated as follows (Hormann et al., 1994; Murchie and Lawson, 2013):

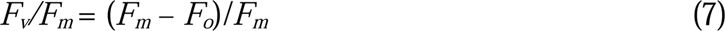

Images of the three plant species at each PPFD were captured. One representative image for each species and each treatment at a PPFD of 189 μmol m^−2^ s^−1^, which was the closest PPFD to the plant growth light condition, was selected and is shown in Figure 5A.

#### 2.2.4. Plant growth analysis

After treatment with LED 664 or LD 660 for 12 d, the plants were harvested, and the shoots (leaves) and roots were separated using sharp scalpels. After the leaf area (cm^2^) was measured, the whole shoot was dried in an oven at 80 °C for 72 h, and the shoot dry weight (mg) was determined. The leaf mass per area (mg cm^−2^) was calculated as the shoot dry weight divided by the leaf area.

### 2.4. Statistical analysis

To ensure the accuracy of the gas exchange statistical analyses and minimize the impact of individual differences, we analyzed the experimental data using a generalized linear mixed model (GLMM). Multiple comparisons of means were subsequently conducted with the Tukey□Kramer honest significant difference (HSD) test (*P* < 0.05) using R software version 4.2.2 (R Development Core Team, 2021). The gas exchange parameter data were visualized through box plots generated with GraphPad Prism 9 (GraphPad Software Inc., San Diego, CA, USA). In these plots, data dispersion was represented by the height of the box, whereas average values were indicated by the lines within the box. The maximum and minimum values are denoted by the highest and lowest points of the vertical line, respectively. The fitted PPFD response curves of the chlorophyll parameters were analyzed using nonlinear regression with GraphPad Prism 9, which employs the asymmetrical (five-parameter) equation. Additionally, comparisons of the data means, excluding gas exchange, were performed with a *t* test (\**P* < 0.05, \*\**P* < 0.01, and \*\*\**P* < 0.001) using SPSS 26.0 statistical software (SPSS Inc., Chicago, IL, USA).

## 3. Results

### 3.1. Effects of different red light spectra on photosynthesis

The gas exchange parameters of tobacco leaves varied significantly with different red light sources (Figure 2). Among the LED and LD lights, the LED 664 and LD 660 lights presented the highest net photosynthetic rates, with the rate of the LD 660 light being 19.1% higher than that of the LED 664 light. In contrast, leaves exposed to LED 629 and LD 635 had significantly lower net photosynthetic rates than those exposed to LD 660 or LED 664. Leaves exposed to LD 673 and LD 690 showed the lowest photosynthetic rates (Figure 2A). For stomatal conductance, tobacco leaves exposed to LD 660 and LED 664 exhibited high values, followed by slightly lower values in leaves irradiated with LED 629 and LD 635. The lowest stomatal conductance was observed in leaves exposed to LD 673 and LD 690 (Figure 2B). The intrinsic water use efficiency (WUE_i_) was generally similar across the different light sources, with the highest efficiency observed under LD 660, followed by LED 664 (Figure 2C).

**Figure 2:**
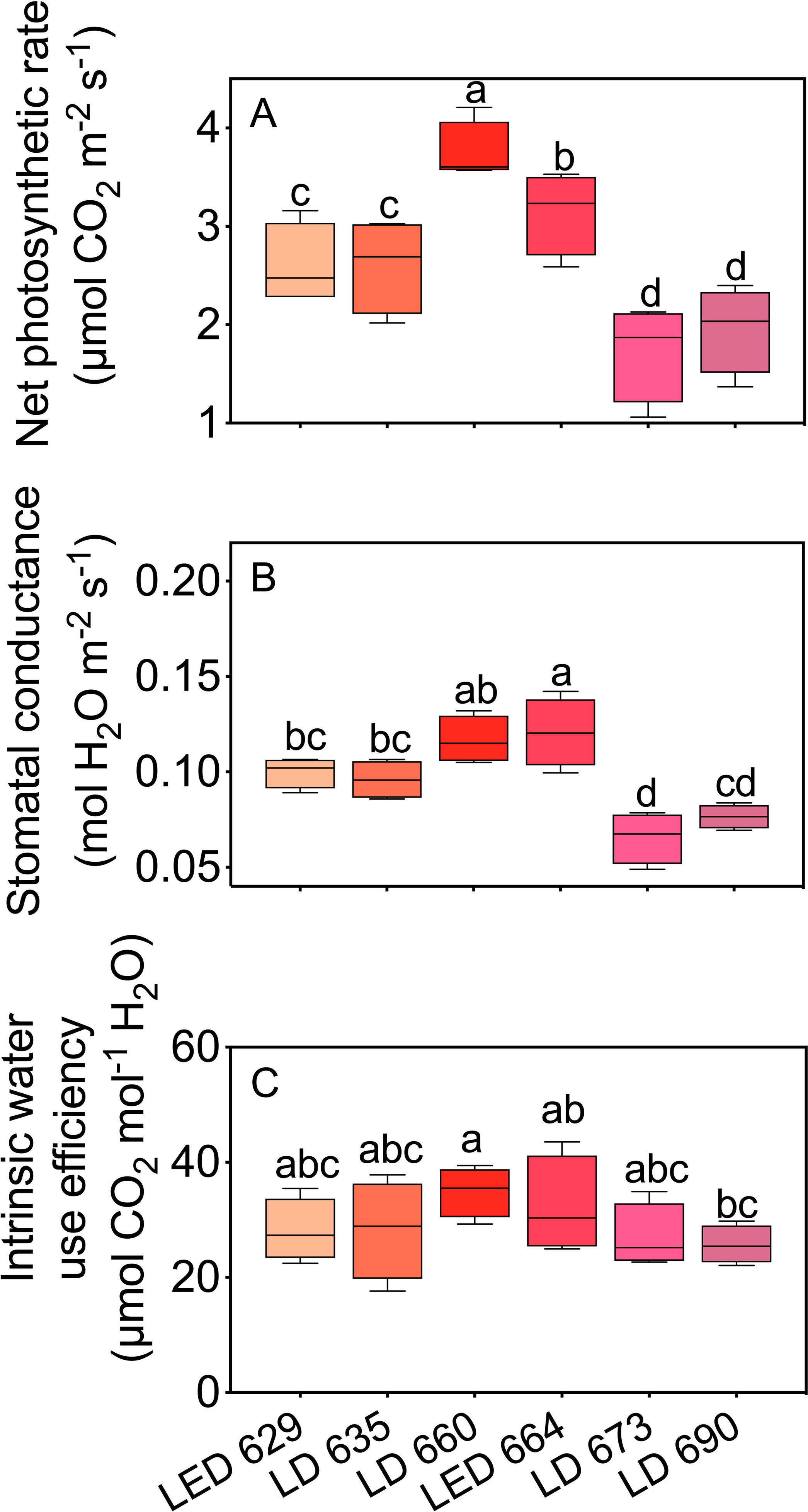
The net photosynthetic rate (A), stomatal conductance (B), and intrinsic water use efficiency (C) of tobacco leaves irradiated with different LED or LD lights. Different letters in each column indicate significant differences at *P* < 0.05, according to the Tukey–Kramer HSD test. The data are presented as the mean ± SE, n = 4.

The leaves of tobacco plants grown under identical environmental conditions were used to test the photochemical efficiency of PS□, represented by Y(□) and NPQ, and the redox state of PS□, represented by qL, when irradiated with LED 664 and LD 660. Compared with those irradiated with LED 664, the leaves irradiated with LD 660 exhibited Y(□) and qL values that were 7.2% and 18.3% higher, respectively (Figure 3). There was no significant difference in NPQ between the two light sources.

**Figure 3:**
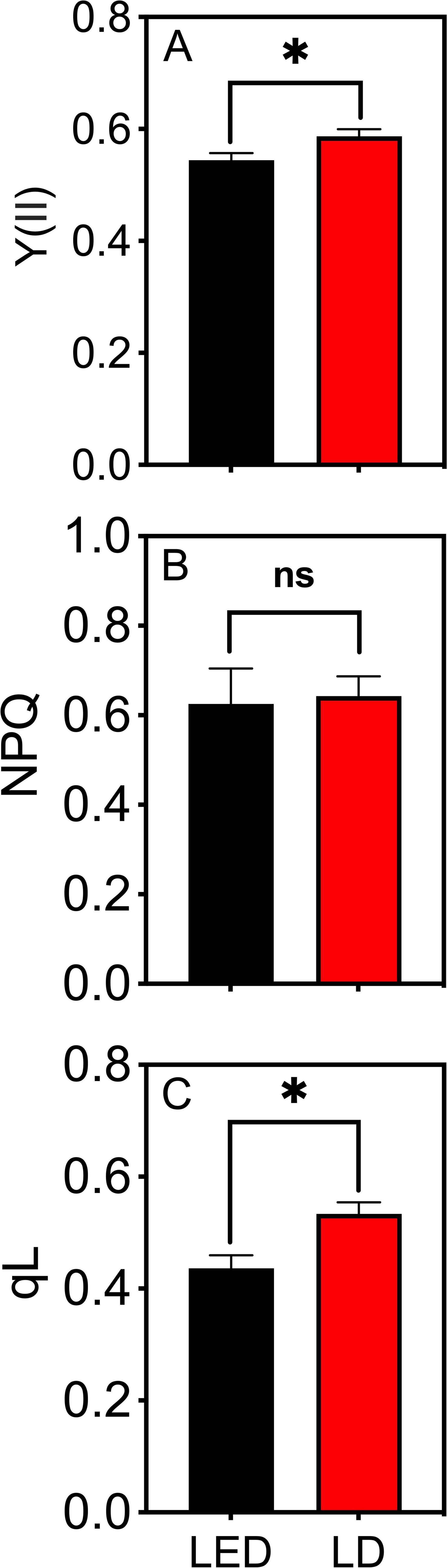
Y(□) (A), NPQ (B), and qL (C) of tobacco leaves irradiated with LED 664 or LD 660 with a PPFD of 150 μmol m^−2^ s^−1^. * indicates a significant difference at *P* < 0.05 according to a *t* test. The data are presented as the mean ± SE, n = 4.

### 3.2. Carbohydrate accumulation

The starch content in plants irradiated with LD 660 for 8 h was 18% higher than that in plants irradiated with LED 664 (Figure 4A). However, the sucrose, glucose, and fructose contents in plants under red LED and LD light were not significantly different.

**Figure 4:**
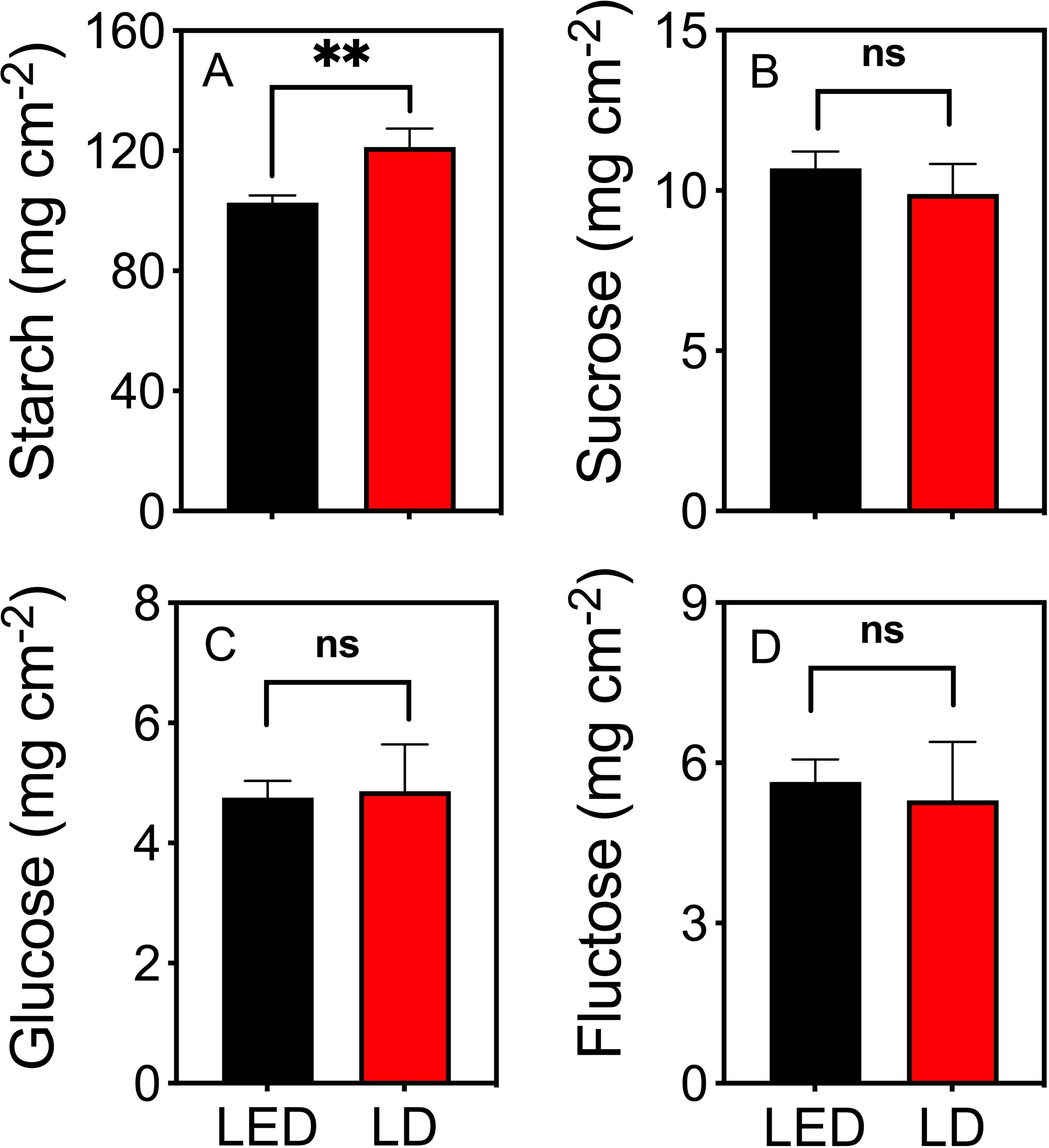
The contents of starch (A), sucrose (B), glucose (C), and fructose (D) in tobacco plants after irradiation with red LED 664 or LD 660 for 8 h. ** indicates a significant difference at *P* < 0.01, ns indicates no significant difference according to the *t* test. The data are presented as the mean ± SE, n = 4.

### 3.3 Spectral characteristics of LED 664 and LD 660 lights

The values of the yield photon flux density (YPFD) and the phytochrome photoequilibria (PPE) for these two light sources were similar (Table 1), although the spectra of LED 664 and LD 660 differed (Figure 1).

**Table 1.**
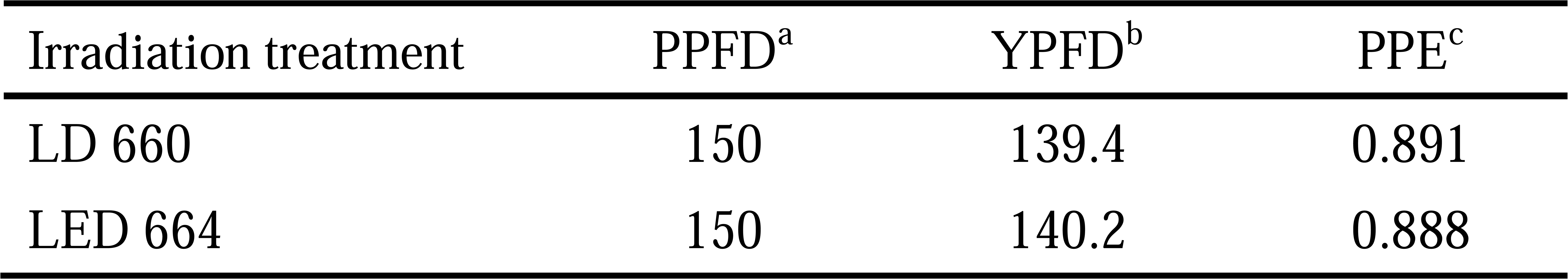
Spectral characteristics of LED 664 and LD 660. ^a^PPFD: Photosynthetic photon flux density, μmol m^−2^ s^−1^. ^b^YPFD: Yield photon flux density, μmol m^−2^ s^−1^, which is the product of the incident flux density of SPD and relative quantum efficiency, based on McCree (1972) and Sager et al. (1988). ^c^PPE: Phytochrome photoequilibria, which is the estimated P_r_/P_total_ following Sager et al. (1988).

### 3.4. Chlorophyll fluorescence parameters at different PPFDs

The photosynthetic efficiency of PSII reaction centers (Y(II)) was examined in tobacco, Arabidopsis, and lettuce plants after being grown under continuous LED 664 or LD 660 for 12 d (Figure 5). The Y(□) value for tobacco plants grown under LD 660 started at 0.75 (F_v_/F_m_) and gradually decreased to 61.3% of the maximum value as the PPFD increased from 0 to 467 μmol m□² s□¹. In contrast, for LED 664, the Y(□) began at 0.61 and sharply declined to 26.2% of the maximum value. Similar trends were observed in Arabidopsis plants with increasing light intensities. The Y(□) values for those grown under LD 660 started at 0.79 and gradually decreased to 54.5% of the maximum value, whereas for those grown under LED 664, Y(□) began at 0.63 and declined to 28.1% of the maximum value. The Y(□) values of tobacco and Arabidopsis plants grown under LD 660 at each PPFD significantly exceeded those grown under LED 664. Conversely, no significant difference was observed between the Y(□) values of lettuce plants grown under LD 660 and those grown under LED 664. The respective Y(□) values of lettuce grown under the LD 660 and LED 664 treatments started at 0.72 and 0.73 and gradually decreased to 0.36 and 0.38 as the PPFD increased (Figure 5B).

**Figure 5:**
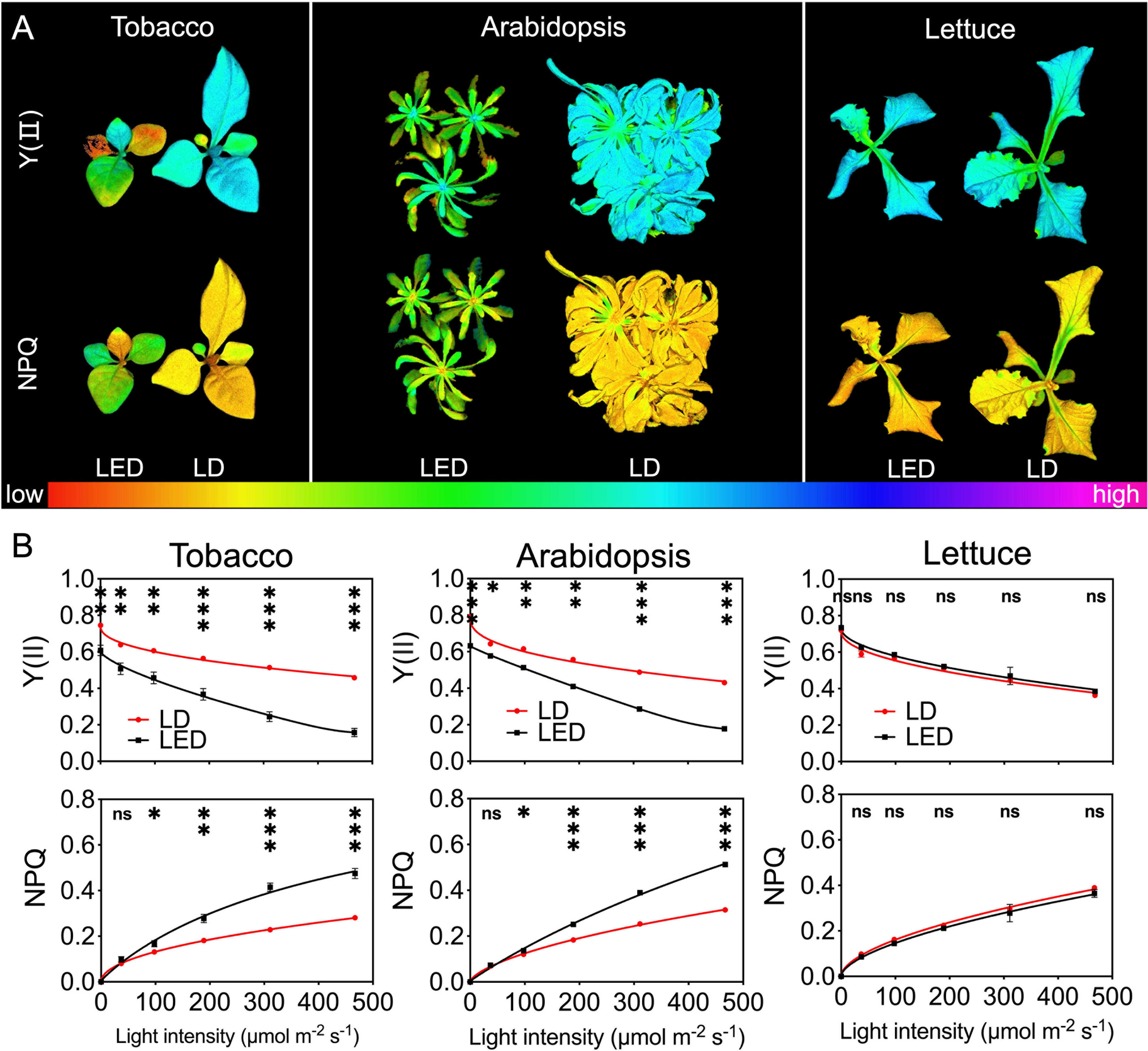
Chlorophyll fluorescence parameters of tobacco, Arabidopsis, and lettuce plants after 12 d of growth under continuous LED 664 or LD 660, with a PPFD of 150 μmol m^−2^ s^−1^. (A) Representative images of Y(□) and NPQ in the three species of plants. (B) Response of Y(□) and NPQ to different PPFD curves of the three species of plants. *** indicates a significant difference at *P* < 0.001, ** indicates a significant difference at *P* < 0.01, * indicates a significant difference at *P* < 0.05, ns indicates no significant difference according to the *t* test. The data are presented as the mean ± SE, n = 4.

All plants exhibited an increase in NPQ values with increasing PPFD. Notably, compared with those grown under LD 660, the NPQ of tobacco and Arabidopsis plants grown under LED 664 increased more rapidly. For tobacco and Arabidopsis, the NPQ value was significantly lower under LD 660 than that under LED 664 when the PPFD exceeded 37 μmol m□² s□¹. In contrast, lettuce plants under both LD 660 and LED 664 exhibited a similar rate of increase in NPQ, with no significant differences observed between LD 660 and LED 664 at any PPFD (Figure 5B).

### 3.5. Biomass accumulation and morphology in plants

Compared with those grown under LED 664, tobacco plants grown under LD 660 displayed a greener color and greater overall size, with larger laminae and longer petioles (Figure 6A). Additionally, compared with those grown under LED 664, the lettuce plants grown under LD 660 presented elongated petioles, increased leaf areas, and a larger overall size (Figure 6A). Moreover, Arabidopsis plants grown under LED 664 exhibited severe chlorosis and anthocyanin pigmentation, along with a smaller overall size compared with plants grown under LD 660 (Figure 6A). Specifically, the shoot dry weights of tobacco, Arabidopsis, and lettuce grown under LD 660 were 1.75, 1.57, and 1.28 times greater, respectively, than that of plants grown under LED 664 (Figure 6B). Additionally, the respective leaf areas of these three plants under the LD 660 treatment were 2.10, 2.28, and 1.70 times greater than those grown under LED 664 (Figure 6B). Conversely, the leaf mass per area of tobacco, Arabidopsis, and lettuce under the LD 660 treatment was 0.84, 0.68, and 0.84 times lower, respectively, than that of plants grown under the LED 664 treatment, (Figure 6B).

**Figure 6:**
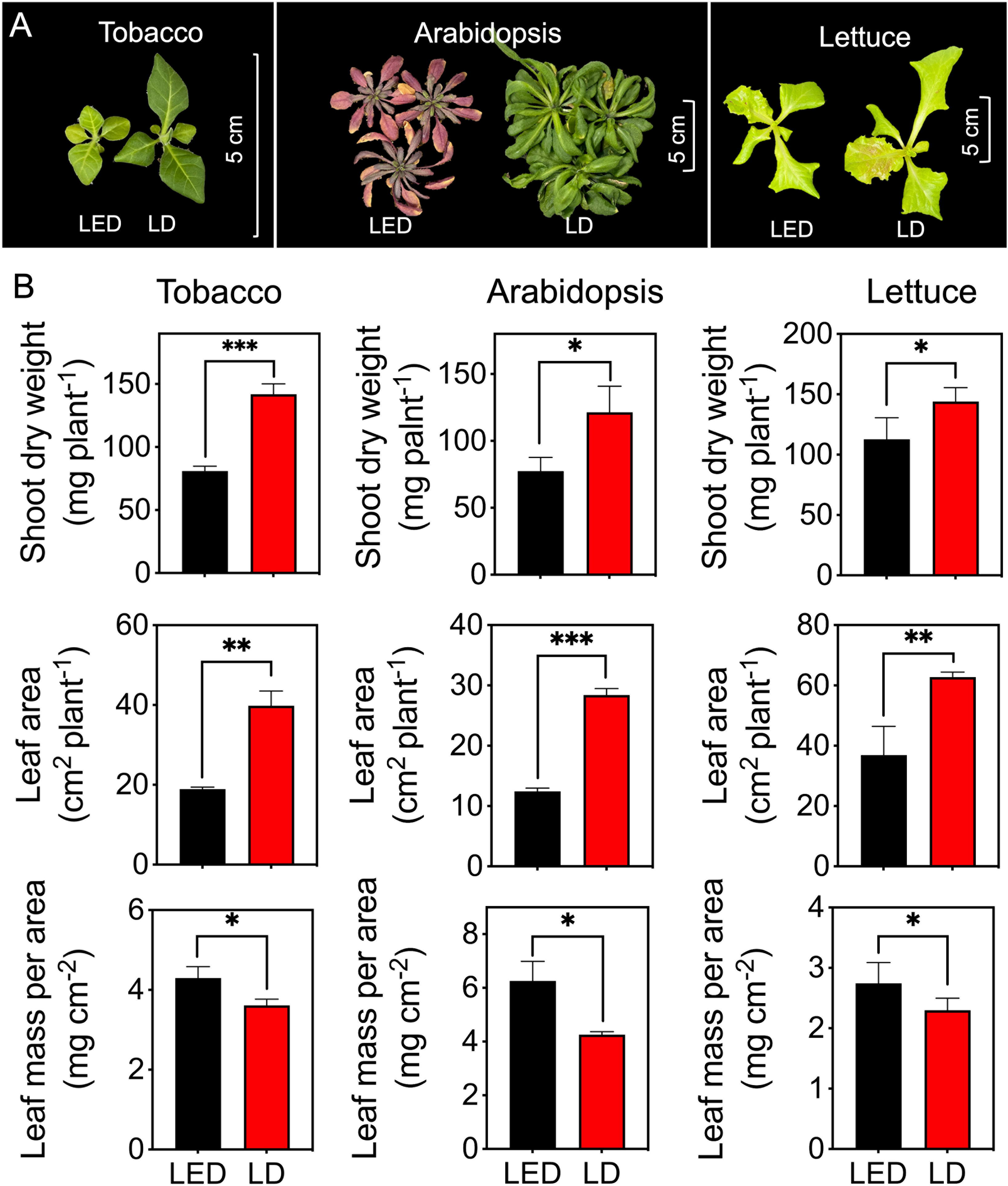
The growth indices of tobacco, Arabidopsis, and lettuce plants after 12 d of growth under continuous LED 664 or LD 660 with a PPFD of 150 μmol m^−2^ s^−1^ for 12 d. (A) Representative images of the three species of plants. (B) The shoot dry weight, leaf area, and leaf mass per area of the three species of plants. *** indicates a significant difference at P < 0.001, ** indicates a significant difference at P < 0.01, * indicates a significant difference at P < 0.05 according to the *t* test. The data are presented as the mean ± SE, n = 4.

## 4. Discussion

To support the growing global population, researchers are increasingly adopting sustainable strategies. Indoor horticulture must also prioritize sustainability. Given that artificial light systems serve as the primary light source in indoor horticulture, ongoing investigations are aiming to optimize artificial lighting systems to support plant growth (Hogewoning et al., 2010; Ohtake et al., 2018; Li et al., 2021). Although traditional light sources and LEDs have been extensively studied (Van Delden et al., 2021), laser diodes (LDs) remain relatively unexplored in the context of photosynthesis, despite their unique advantages. The present study clearly demonstrates that LDs enhanced photosynthesis and plant growth compared with LEDs with similar peak wavelengths, suggesting that LDs offer significant advantages over LEDs for indoor plant production.

### 4.1. Fine-tuning red light to explore wavelength-specific impacts on photosynthetic efficiency

Red light plays a crucial role in regulating plant growth and development because it is significantly absorbed by chlorophyll (Runkle, 2016). Previously, McCree (1972) demonstrated that red light with emission peaks at 600 or 625 nm provides the highest quantum yield for CO_2_ assimilation (mol CO_2_ assimilated per mol photons absorbed) across 22 plant species. Subsequent analysis by Inada (1976) identified 625, 650, and 675 nm as the optimal wavelengths for photosynthetic efficiency in red light. In this study, we demonstrated that the net photosynthetic rates were ranked as follows: LD 660 > LED 664 > LED 629 ≈ LD 635 > LD 673 ≈ LD 690, where “>” indicates a significant difference and “≈” indicates no significant difference (Figures 1, 2A). Furthermore, LD 660 and LED 664 exhibited relatively high stomatal conductance, whereas LD 673 and LD 690 showed the lowest values, which aligns with trends in the net photosynthetic rate across the six light sources (Figures 2A and 2B).

These findings suggest that red light promotes stomatal opening by reducing intercellular CO_2_ (*C*i) through red light-driven mesophyll photosynthesis (Shimazaki et al., 2007). However, the differences in stomatal conductance across treatments were less pronounced than the differences in net photosynthetic rates, implying that additional mechanisms influence stomatal behavior. For example, Messinger et al. (2006) reported that red light can stimulate stomatal opening even when *C*_i_ remains constant, whereas Kromdijk et al. (2019) and Taylor et al. (2024) proposed that red light may modulate the redox state of plastoquinone (PQ), potentially signaling beyond the chloroplast to regulate stomatal responses. In addition, Ando and Kinoshita (2018) suggested that red light can promote stomatal opening by inducing H+-ATPase phosphorylation in guard cells. These mechanisms, while promising, remain incompletely understood. Our results indicate that variations in red light wavelength and spectral width influence stomatal responses in ways that do not consistently align with net photosynthetic rates. Interestingly, LD 660 exhibited high net photosynthetic rate and stomatal conductance but maintained a WUE_i_ comparable to that of other treatments (Figure 2C). This finding indicates that LD 660 achieves efficient CO_2_ assimilation while conserving water (Lawson and Blatt, 2014).

This study provides nuanced insights into how specific red-light wavelengths affect photosynthesis, stomatal conductance, and WUE_i_. The superior performance of LD 660 and LED 664 highlights the importance of fine-tuning light spectra to optimize both photosynthetic efficiency and WUE_i_, in contrast to earlier studies emphasizing broader wavelength ranges (Klimek-Kopyra et al., 2021). These findings underscore the necessity for precision in controlled-environment agriculture, where spectral specificity plays a crucial role in optimizing photosynthetic performance. Overall, our results emphasize that red light with distinct spectral characteristics differentially influences photosynthetic processes and associated physiological responses.

### 4.2. Optimizing photosynthesis with narrowband red LD light for increased plant growth and yield

Our study found that the net photosynthetic rate of plants under LD 660, which has a narrow waveband (< 10 nm), was significantly higher than that under LED 664, which has a broader waveband (~50 nm), despite both light sources having identical PPFD levels and similar emission peak positions (Figures 1 and 2). This finding indicates that, even when spectra share similar peaks, differences in waveband width can significantly affect photosynthetic efficiency. Because photosynthesis underpins plant growth and yield (Taiz and Zeiger, 2015; Yamori et al. 2016), the increased photosynthetic efficiency observed under red LD light likely plays a key role in promoting plant growth (Figure 5B). Furthermore, carbohydrates, the end products of CO_2_ assimilation during photosynthesis, accumulated at higher levels under red LD light than under LED light (Figure 4A). These findings highlight the effectiveness of red LD in promoting plant photosynthesis and yield compared with red LEDs, even with a comparable yield photon flux density (YPFD) (Table 1).

In addition to photosynthesis, plant morphology is crucial for the accumulation of dry matter (Li et al., 2021). In this study, plants exposed to red LD light developed a significantly larger total leaf area compared with that of plants exposed to LED light (Figure 5B). We hypothesized that the observed morphology might reflect characteristics of shade-avoidance syndrome (SAS), which is typically induced by a low red-to-far-red (R:FR) light ratio sensed by phytochromes. SAS is characterized by traits such as stem and petiole elongation and increased leaf expansion (Park and Runkle, 2017; Legris et al., 2019). However, calculations of the phytochrome photoequilibria (PPE), an estimate of the P_r_/P_total_ ratio (Sager et al., 1988), yielded nearly identical values for LD 660 (0.891) and LED 664 (0.880) (Table 1). These findings suggest that the morphological differences between plants grown under LD 660 and LED 664 are not due to variation in far-red light absorption by phytochromes.

The redox balance between PSI and PSII may also influence plant physiological responses (Foyer and Noctor 2005; Huner et al., 1998). Under identical growth conditions, the qL parameter of PSII was 18.3% higher in plants exposed to red LD light than that in plants exposed to LED light, with Y(II) being 7.2% higher (Figure 3). This significant increase in qL suggests that LD light may increase the electron transport efficiency of PSII by more effectively activating PSI. Enhanced PSI activity likely facilitates more efficient electron flow, reducing bottlenecks in the electron transport chain and allowing more PSII reaction centers to remain open. As PSI is more frequently excited, it drives greater electron transfer toward NADP^+^, maintaining the electron transport system in a more oxidized state (Kramer et al., 2004; Nelson and Yocum, 2006). This oxidized state creates a signaling environment that enables plant acclimation. For example, under high temperatures and elevated CO_2_ concentrations, plants may develop “shade-type” characteristics such as thinner and larger leaves (Way and Yamori, 2014; Leakey et al., 2009). These adaptations optimize the leaf area for effective light and CO_2_ capture, reduce ROS accumulation, and prevent photoinhibition. In this study, the significantly increased leaf area and decreased leaf mass per area observed in tobacco, Arabidopsis, and lettuce plants grown under red LD light compared with those grown under LED light (Figure 5B) may be linked to changes in redox states within plant cells. The increased leaf area likely contributed to the increased shoot dry weight observed in these plants.

Based on our findings, LDs demonstrate remarkable potential for advancing agricultural practices. Their ability to optimize photosynthesis and support beneficial plant morphology positions them as valuable tools for future agricultural technology. Adopting LD-based systems could lead to higher yields, improved crop quality, and more sustainable food production practices.

### 4.3. Impact of continuous red light on photosynthesis and plant tolerance across species

Continuous light is often considered a strategy to increase plant productivity by extending the photoperiod, allowing photosynthetic organisms to assimilate more CO_2_ daily (Kalaitzoglou et al., 2019; Xu et al., 2021; Proietti et al., 2021). To evaluate this potential, we tested the effects of continuous LD 660 compared with continuous LED 664 on plant production. However, continuous light is also known to induce negative effects on plant growth and physiology (Velez-Ramirez et al., 2011; Demers et al., 1998). Consistent with these findings, tobacco and Arabidopsis plants exposed to continuous red LED light for 12 d presented reduced photosynthetic capacity (Figure 6). Our data showed a significant decline in F_v_/F_m_, with values falling below the acceptable range for normal plant growth (Murchie and Lawson, 2013). Additionally, as PPFD increased, Y(II) levels decreased sharply (Figure 6B), indicating photoinhibition of PSII and reduced photochemical efficiency. This likely impaired carbon fixation during photosynthesis (Demmig et al., 1987; Dodd et al., 1998).

The increased nonphotochemical quenching (NPQ) values across light intensities further support the notion that continuous red LED light resulted in inefficient light energy utilization (Figure 6B). Previous studies have reported that continuous light negatively affects photosynthetic performance by reducing photosynthetic capacity, quantum yield, Rubisco carboxylation rates, and electron transport efficiency (Murage et al., 1997; Kang et al., 2013; Zha et al., 2019a; Heyneke et al., 2013; Van Gestel et al., 2005). These effects are often attributed to excess light energy, leading to the accumulation of ROS and the accumulation of starch and sugars (Golan et al., 2006; Heyneke et al., 2013; Van Gestel et al., 2005; Velez-Ramirez et al., 2011). These factors inhibit photosynthesis and contribute to chlorosis and tissue degradation, which were also observed in our study (Figure 5A).

The plants exposed to continuous red LED light also developed higher leaf mass per area (Figure 5B), suggesting structural adjustment to mitigate photoinhibition. This aligns with prior research indicating that plants acclimate to continuous light by thickening their leaves to dissipate excess energy and protect the photosynthetic apparatus (Arve et al., 2013; Zha et al., 2019b). Such acclimation may also reduce water loss and improve light utilization efficiency. In contrast, tobacco and Arabidopsis plants exposed to continuous red LD light exhibited greater tolerance to continuous light. This was evidenced by higher F_v_/F_m_ values, increased Y(II) values across light intensities, lower NPQ values at light intensities exceeding 37 μmol m□² s□¹, and reduced chlorosis compared with those of plants under red LED light (Figures 5A and 6). These results are consistent with studies demonstrating the positive effects of laser light on plant photosynthesis. For example, Ooi et al. (2016) reported that compared with cool-white fluorescent light, continuous red and blue laser light reduced the expression of the light stress marker genes *APX1* and *GST6*. Additionally, laser light has been shown to increase resistance to environmental stressors, such as salinity, drought, and oxidative stress, by modulating gene expression and increasing antioxidant enzyme activity, photosynthesis, and nutrient uptake (Ali et al., 2020; Qiu et al., 2013; Gao et al., 2015).

Interestingly, lettuce showed greater tolerance to continuous light than did tobacco and Arabidopsis, with no significant differences in chlorophyll fluorescence parameters between the continuous red LED and LD light treatments (Figure 6B). Furthermore, the lettuce did not exhibit any observable signs of continuous light-induced injury (Figure 6A). These results align with those of Zha et al. (2019a), who found that lettuce could maintain normal growth under continuous light at a PPFD of 200 μmol m□² s□¹ for 15 d, despite elevated ROS and antioxidant levels.

Environmental factors, such as the PPFD, can further influence plant responses to continuous light. Higher PPFD intensities exacerbate continuous light-induced damage, as shown by reductions in F_v_/F_m_ and quantum yield in lettuce grown under continuous light with a PPFD of 300 μmol m□² s□¹ (Zha et al., 2019b). In a supplementary experiment, lettuce plants exposed to continuous red LED and LD light at a PPFD of 300 μmol m□² s□¹ exhibited a slightly lower F_v_/F_m_ than that of plants grown at 150 μmol m□² s□¹. However, no significant differences were observed between the red LED and LD treatments in terms of chlorophyll fluorescence parameters (Supplemental Figure 1B). Notably, the anthocyanin content was significantly higher in lettuce under red LED light than that under red LD light at 300 μmol m□² s□¹ (Supplemental Figure 1C), indicating light stress. Anthocyanin synthesis is often triggered when the amount of radiation absorbed exceeds the capacity of the photosynthetic system (Smillie and Hetherington, 1999; Trojak and Skowron, 2017). Thus, the greater anthocyanin accumulation under continuous red LED light suggests a stress response greater than that under red LD light.

## Conclusions

Our findings demonstrate that laser diode (LD) light, particularly with a peak at 660 nm, offers distinct advantages over LED light for plant growth in indoor horticulture. The plants grown under LD 660 exhibited higher photosynthetic efficiency, increased carbohydrate accumulation, and greater tolerance to continuous light as well as improved morphological traits, including an expanded leaf area and increased biomass production. These benefits highlight the potential of LDs for optimizing resource use and improving adaptability in controlled-environment agriculture. The adoption of LD technology could contribute to sustainable urban agriculture and address the rising demand for food in an increasingly constrained agricultural landscape. Further research is needed to optimize LD applications and elucidate their underlying beneficial effects on plant physiology.

## CRediT authorship contribution statement

Lie Li: Data curation, Investigation, Formal analysis, Writing – original draft. Ryusei Sugita: Conceptualization, Investigation, Resources. Kampei Yamaguchi: Conceptualization, Resources. Hiroyuki Togawa: Conceptualization, Resources. Ichiro Terashim: Writing – review and editing. Wataru Yamori: Conceptualization, Investigation, Funding acquisition, Writing – review and editing.

## Data statement

Supporting data can be requested by contacting the corresponding author.

## Figure captions

Supplemental Figure 1: Lettuce plants were grown under continuous LED 664 or LD 660 with a PPFD of 300 μmol m^−2^ s^−1^ for 12 d. (A) Representative image of a lettuce plant. (B) Response of the photosynthetic fluorescence parameters (Y(□) and NPQ) to different light intensities. (C) Accumulation of anthocyanin in the lettuce plants. ** indicates a significant difference at *P* < 0.01, ns indicates no significant difference according to the *t* test. The data are presented as the mean ± SE, n = 4.

