## Supplementary figures and images for "High-Precision Lighting for Plants: Laser Diodes Outperform LEDs in Photosynthesis and Plant Growth"

### Supplemental Figure

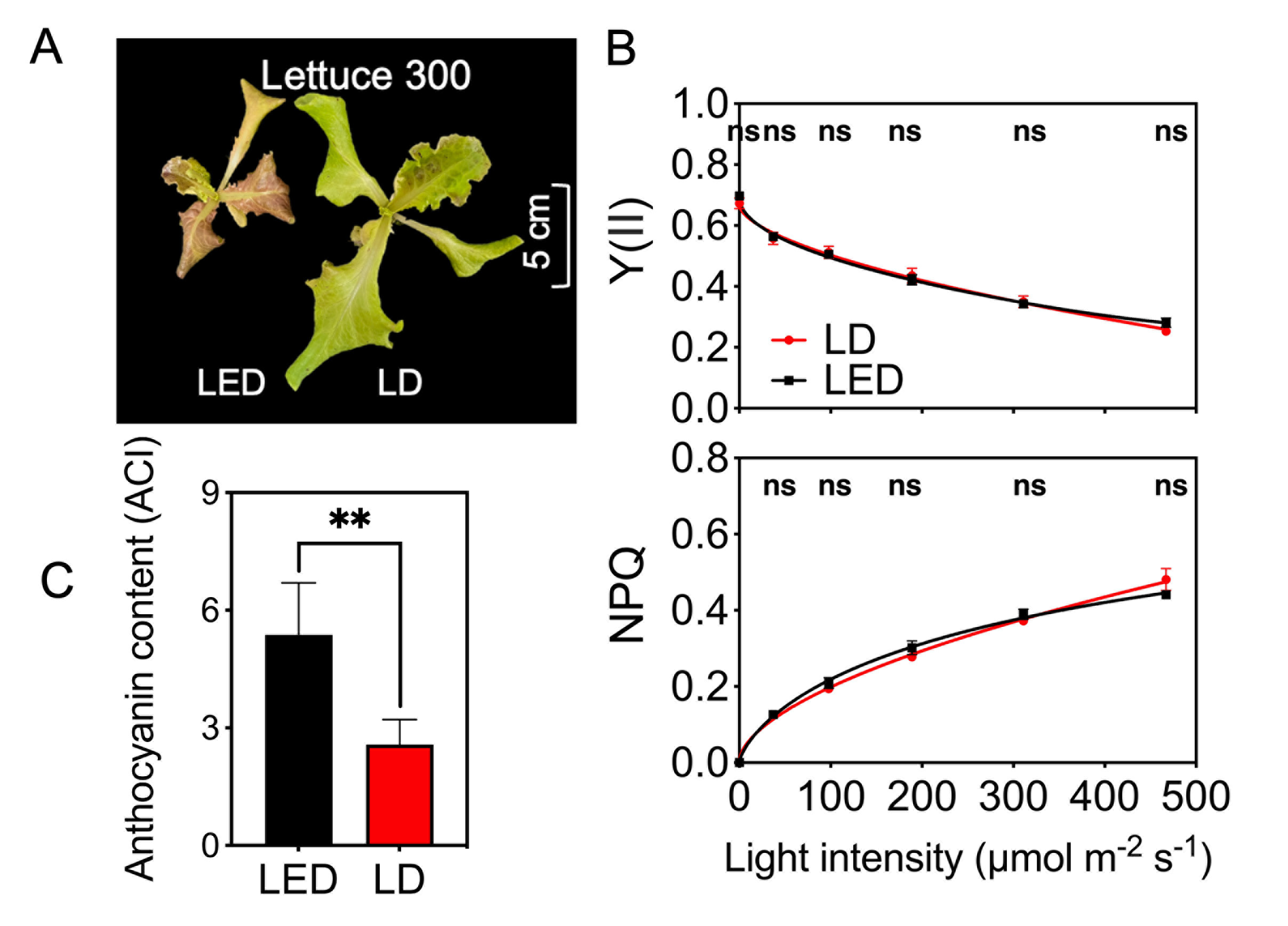
